# AmesNet: A Task-Conditioned Deep Learning Model with Enhanced Sensitivity and Generalization in Ames Mutagenicity Prediction

**DOI:** 10.1101/2025.03.20.644379

**Authors:** Tyler Umansky, Virgil Woods, Sean M. Russell, Daniel Haders

## Abstract

Regulatory agencies require comprehensive genotoxicity assessments for all novel small-molecule therapeutics prior to human trials. Developers often delay these studies until a candidate is nearing regulatory submission because they are expensive and secondary to bioactivity. This timing creates a bottleneck where late-stage failures can jeopardize >$10 million in capital and multiple years of developmental progress per candidate. The Ames assay is used to detect a molecule’s mutagenic potential. Regulators now explicitly support the use of *in silico* Ames mutagenicity models through enabling legislation, dedicated FDA AI toxicology programs, internationally harmonized guidelines, and benchmark challenges. However, current Ames models suffer from a dramatic sensitivity drop-off when they evaluate molecules outside their training domain. Sensitivity is the most important metric in Ames prediction because false negatives allow mutagenic compounds to advance undetected and trigger the most costly late-stage failures. Attempts to fix this sensitivity drop-off often reduce overall model performance, which can be represented by balanced accuracy. For example, DeepAmes reports high levels of sensitivity only by sacrificing its balanced accuracy. We introduce AmesNet, a novel Task-Conditioned modeling paradigm that achieves both class-leading sensitivity and balanced accuracy in novel chemical spaces. AmesNet utilizes a dual branch architecture containing a molecular encoder and a dedicated channel to condition Ames assay context such as metabolic activation and bacterial strain type. In comparative benchmarks, AmesNet reached a sensitivity of 0.73 (95% confidence interval: 0.68–0.77) and a simultaneous balanced accuracy of 0.81 (95% confidence interval: 0.79–0.83) on the out-of-domain test data. This represents an improvement in sensitivity of up to 46% over existing approaches without a trade-off in balanced accuracy. Structural analysis demonstrates that AmesNet recovers difficult-to-detect mutagenic compounds missed by existing models. This framework provides a high-confidence filtering mechanism that enables drug developers to turn a costly late-stage safety bottleneck into a proactive decision-making edge.

## Introduction

New small-molecule pharmaceuticals must undergo a suite of toxicity testing before entering clinical trials. The FDA and international regulators expect Ames genotoxicity data in regulatory submissions as described in the ICH S2(R1) guideline[1–3]. The Ames test under Good Laboratory Practice (GLP) conditions exceeds $10,000 per compound. This cost excludes the chemical sample price[4]. Such high expenditure makes routine screening of large compound libraries impractical during early drug discovery. GLP Ames testing typically occurs near the regulatory submission stage after significant investment into the preclinical candidate[5]. Preparing for human clinical trials takes approximately 3 years in the average small-molecule discovery timeline. This process can require >$10 million per candidate[6]. An Ames test failure at this late stage results in years of wasted effort and millions of dollars in lost capital.

In response to such bottlenecks, the FDA and regulatory agencies around the world have formalized *in silico* approaches as a pathway to reduce and outright replace traditional preclinical testing. The FDA Modernization Act now provides the legal framework for AI-based computer models to support and replace wet-lab and animal testing for drug evaluation[7]. Accordingly, the FDA established programs like the Artificial Intelligence Program for Toxicology (AI4TOX) to operationalize AI within safety reviews.[8] Under this program the SafetAI initiative focuses on developing AI models for toxicological endpoints. DeepAmes was developed in this initiative to predict Ames mutagenicity for use within regulatory frameworks[9, 10]. Internationally, the ICH M7(R1) guideline explicitly endorses Quantitative Structure-Activity Relationship (QSAR) methodologies to predict Ames mutagenicity[11]. The First and Second Ames/QSAR International Challenge Projects further propelled regulatory acceptance. During these projects, government agencies in the U.S., Japan, and Italy used standardized benchmarks to improve model performance for Ames prediction[12, 13].

The Ames test uses specialized bacterial strains in growth-restrictive conditions to detect if a drug candidate might cause genetic mutations. Mutagenic potential is evidenced by bacterial growth after exposure to either the parent compound or its metabolic byproducts (Figure 1A)[1, 2, 14]. Effective screening therefore depends on high sensitivity to avoid misclassifying a dangerous molecule as safe. Meeting this requirement remains a challenge for Ames QSAR models. Low sensitivity is a critical failure because it leads to more genotoxic compounds advancing undetected[12, 13]. Such failures are amplified when AI models encounter novel chemical structures distinct from their training data. This challenge in novel chemical space is known as out-of-domain (OOD) generalization. Achieving operational Ames model utility simultaneously requires high sensitivity and OOD generalization so that no genotoxic compound escapes detection when screening novel chemical libraries[12, 13]. Benchmark data from the Second Ames/QSAR International Challenge Projects revealed a significant collapse in sensitivity for current Ames models when encountering novel chemotypes[13]. The project yielded a participant average sensitivity of only 0.46 on the OOD test set. High-profile models like the FDA’s DeepAmes achieved a sensitivity of 0.47 and MIT’s ChemProp yielded 0.32[13]. Attempts to increase sensitivity have forced trade-offs in overall model performance, which can be measured by balanced accuracy. DeepAmes more recently reports a sensitivity of 0.87 with a corresponding balanced accuracy of 0.52, providing little predictive signal[9, 10]. These results demonstrate that current methods cannot maintain the sensitivity required for the reliable safety screening of novel drug candidates. Such unreliability creates a pathway for unidentified genotoxic molecules to advance into later stage drug development and potentially waste years and millions of dollars in resources.

**Figure 1.**
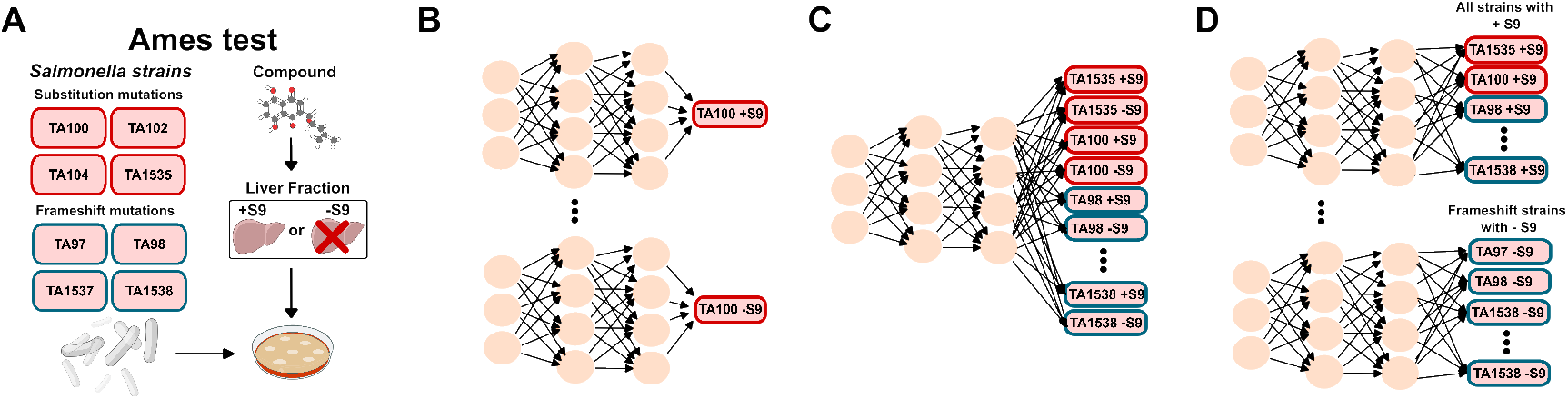
Ames test and predictive algorithms. A) Diagram depicting the Ames test, where the rectangles represent eight strains commonly used. Compounds are tested in the presence or absence of a liver fraction (*±*S9), which mimics metabolic activation in the human body. B) Single-task Ames QSAR algorithm (STL) trained to predict a strain-specific feature, with or without metabolic activation (*±*S9). C) Ungrouped multitask Ames QSAR algorithm (uMTL), trained to predict all strain-specific features, with or without metabolic activation (*±*S9). D) Grouped multitasked Ames QSAR algorithm (gMTL), trained to predict subsets such as substitution strains-specific features with or without metabolic activation (*±*S9)[19].

The pervasive sensitivity gap across current Ames QSAR models necessitates a new modeling framework for maintaining robust performance in novel chemical space. To our best knowledge, all existing AI models for Ames prediction fall under the archetype of Unconditioned Modeling. In this archetype, predictions are generated without the use of an input conditioning channel for explicit assay states like bacterial strain or metabolic activation.

We introduce AmesNet as a novel Task-Conditioned Learning (TCL) paradigm driven by explicit input-based conditioning that diverges from established methods in Ames QSAR modeling. AmesNet predicts strain-specific Ames mutagenicity from the concatenation of an assay state conditioning channel and a molecular encoder branch. AmesNet utilizes an adaptation of ChemPrint for the molecular encoder (Figure 2). ChemPrint is an experimentally validated geometric convolutional neural network within the Model Medicines’ GALILEO platform. The model has documented success across oncology and antiviral programs. These successes include on-target hit rates up to 100%, selectivity improvements exceeding 15,000-fold, and the discovery of MDL-001. MDL-001 has achieved preclinical proof-of-concept across several indications[15– 18]. AmesNet is designed to leverage ChemPrint’s OOD generalization and power this first instantiation of our TCL paradigm for Ames prediction. We propose this framework to close the existing performance gap of Ames QSAR models and enable high-confidence mutagenicity screening in novel chemical space.

**Figure 2.**
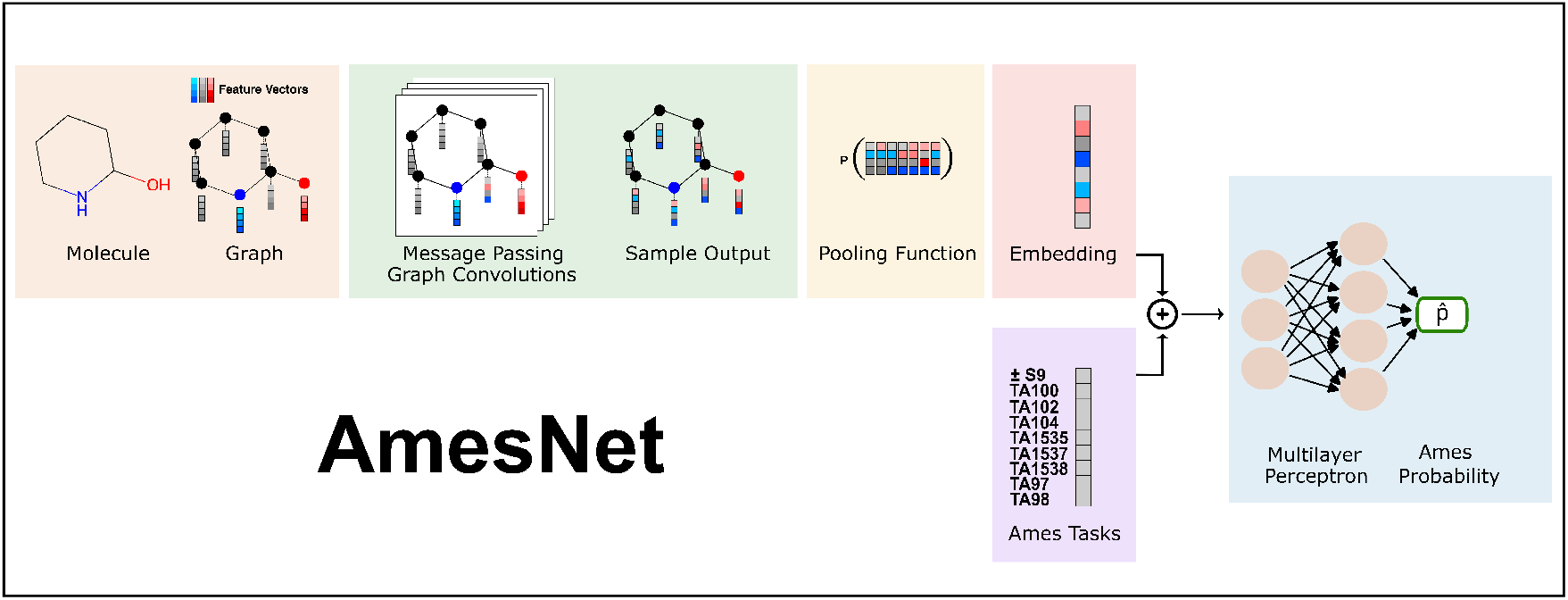
The AmesNet TCL architecture. AmesNet is a dual branch architecture containing a graph convolutional neural network and an Ames task conditioning channel conditioned on strain and *±*S9.

We execute a two-tiered benchmarking strategy to assess AmesNet’s overall performance and to isolate the contribution of TCL for Ames prediction. First, we measure AmesNet’s end-to-end performance relative to established learning frameworks: Single-Task Learning (STL), Ungrouped Multitask Learning (uMTL), and Grouped Multitask Learning (gMTL) (Figure 1B–D)[19]. In this benchmark, we include STL instantiations of ChemProp, GROVER, and DeepAmes[9, 10, 20, 21]. This comparison quantifies AmesNet’s overall performance advantage relative to these comparators. Second, we isolate the contribution of the TCL framework by augmenting established encoders with the conditioning channel: message-passing networks and graph transformers. In this benchmark, we include TCL instantiations of ChemProp and GROVER. This comparison quantifies the performance gains attributable to TCL. We assess performance for each benchmark through the concurrent evaluation of generalization and classification metrics on a withheld OOD test set. Sensitivity is reported as the primary metric. Additionally, we utilize balanced accuracy (BA) and confusion matrices to verify the model’s overall predictive robustness. Across all benchmarks, AmesNet demonstrates substantial improvements in sensitivity, BA, and OOD generalization, outperforming both the Unconditioned Models in the first benchmark and the encoder-swaps in the second benchmark. These results demonstrate that AmesNet and its TCL framework offer enhanced predictive performance in novel chemical space over alternatives.

The performance of AmesNet substantially addresses the OOD sensitivity gap that has limited the utility of Ames QSAR models. AmesNet provides a more reliable pathway for higher-confidence safety screening at the earliest stages of discovery. This framework offers a notably improved alternative to current *in silico* approaches and serves to mitigate the $10 million, three-year risk of late-stage failure due to undetected genotoxicity. This TCL paradigm supports higher-confidence screening of massive chemical libraries and has the potential to transform a late-stage safety liability into a primary strategic advantage.

## Methods

### Data

#### Dataset Source

We attempted to obtain the dataset from the Ames/QSAR International Challenge Projects by reaching out to the authors multiple times but did not receive a response[12, 13]. As a result, we used the best available publicly accessible dataset for this study[19].

The dataset was compiled from four individual data sources: ISSSTY, OASIS, EFSAP, and MHLW. The dataset included Ames test annotations from eight Salmonella strains, each corresponding to a specific mutation type. The base-pair substitution strains are TA100, TA102, TA104, and TA1535. The frameshift mutation strains are TA1537, TA1538, TA97, and TA98. Assays were conducted in the presence or absence of S9 fraction.

#### Dataset Cleaning

The molecules in the dataset were converted from SDF to SMILES format and standardized with respect to charge, stereochemistry, and tautomerism. The dataset was prepared for model training, validation, and testing according to the predefined OOD data splits in the comparator publication[19]. All (compound, strain, S9) triplets with conflicting Ames endpoints were removed from the dataset. Duplicate triplets sharing identical endpoints were collapsed to a single instance. Data leakage across dataset splits was identified and eliminated. Compounds were filtered by molecular weight [100, 1000] Da to retain small molecules only. These filtering steps reduced the training/validation set from 43,897 to 40,129 data points and the test set from 4,838 to 4,528 data points.

#### Dataset Featurization

Each compound is represented as a geometric molecular graph. Nodes correspond to atoms, and edges correspond to chemical bonds. This representation captures the compound’s structural information. Atomic features are encoded as one-hot vectors and provided as node attributes.

Nine one-hot encoded context features represent the Ames assay conditions and are provided through the state conditioning channel. The experimental conditions consist of the Salmonella strain and the presence or absence of S9 metabolic activation. Each data point corresponds to a (compound, strain, S9) triplet with a binary ground-truth Ames label (+/−).

### AmesNet Architecture

#### Task-Conditioned Dual Branch

AmesNet consists of two input branches. The first branch encodes molecular structure using an atomic-level graph encoder that produces a 512-dimensional molecular embedding. The second branch encodes nine one-hot Ames assay context features specifying the Salmonella strain and the presence or absence of S9 metabolic activation. The molecular embedding and context feature vectors are concatenated and passed to a multilayer perceptron for prediction. The molecular branch captures structural information, while the context branch enables conditional mutagenicity prediction. All AmesNet parameters are learned end-to-end from random initialization, and no weights are imported or frozen.

### Model Development Operations

#### Hyperparameter Optimization

Grid search was used to optimize hyperparameters during model training and validation.

#### Compute Resources

All model training and inference were performed on an NVIDIA A100 40GB GPU.

### Reference Models

#### Unconditioned Models

AmesNet was benchmarked against three unconditioned Ames QSAR frameworks: single-task learning (STL), ungrouped multitask learning (uMTL), and grouped multitask learning (gMTL). The multilayer perceptron (MLP) architectures for STL, uMTL, and gMTL follow the reference implementation[19]. In addition, three architectures (DeepAmes, ChemProp, and GROVER) were evaluated as STL baselines by training 16 task-specific models per architecture (bacterial strain x *±*S9)[9, 10, 20, 21]. Each model was trained end-to-end.

#### Task-Conditioned Encoder-Swap Models

AmesNet was also benchmarked by injecting the task conditioning channel into ChemProp and GROVER_large_[20, 21]. Each encoder-swap model was trained end-to-end.

### Metrics and Bootstrapping

Multi-task binary classification performance was evaluated across 16 Ames tasks. Each task was defined as a unique combination of bacterial strain and *±*S9. Sensitivity and balanced accuracy were computed per task and are the principal metrics reported in the main text. Specificity and Matthews correlation coefficient (MCC) were also computed per task and are reported in the Supplementary Information. Each metric was aggregated using a sample-size weighted average, with weights proportional to the number of observations per task. Uncertainty was estimated with a within-task stratified bootstrap (n = 1,000). Positives and negatives were resampled with replacement while also retaining the same task size and class counts. Metrics were recomputed per task and aggregated using the same weights to yield one bootstrap replicate of the estimator. The 95% confidence interval was defined by the 2.5th and 97.5th percentiles of the bootstrap distribution.

## Results

### Benchmark I: AmesNet vs. Unconditioned Modeling Paradigms

#### Benchmark I: Sensitivity on OOD Test Set

Benchmark I evaluated the performance of AmesNet against established Unconditioned Ames QSAR frameworks (STL, uMTL, and gMTL) on sensitivity in an OOD setting. OOD sensitivity and 95% confidence intervals were estimated as described in Methods (Figure 3). AmesNet achieved a sensitivity of 0.73 (95% CI: 0.68–0.77), the highest reported among all models evaluated in Benchmark I. Unconditioned Models showed consistently lower sensitivity. Within the STL framework, the STL-MLP achieved 0.50 (95% CI: 0.45– 0.55), STL-ChemProp achieved 0.74 0.54 (95% CI: 0.49–0.59), STL-GROVER achieved 0.55 (95% CI: 0.50–0.60), and STL-DeepAmes achieved 0.67 (95% CI: 0.62–0.71). Within the uMTL framework, the uMTL-MLP achieved 0.57 (95% CI: 0.52–0.62). Within the gMTL framework, the gMTL-MLP achieved 0.59 (95% CI: 0.55–0.63). AmesNet achieved sensitivity improvements of +0.23 (46%) over STL-MLP, +0.19 (35%) over STL-ChemProp, +0.18 (33%) over STL-GROVER, +0.06 (9%) over STL-DeepAmes, +0.16 (28%) over uMTL-MLP, and +0.14 (24%) over gMTL-MLP. AmesNet exhibits non-overlapping 95% confidence intervals across sensitivity measurements relative to STL-GROVER, STL-ChemProp, STL-MLP, uMTL-MLP, and gMTL-MLP baselines.

**Figure 3.**
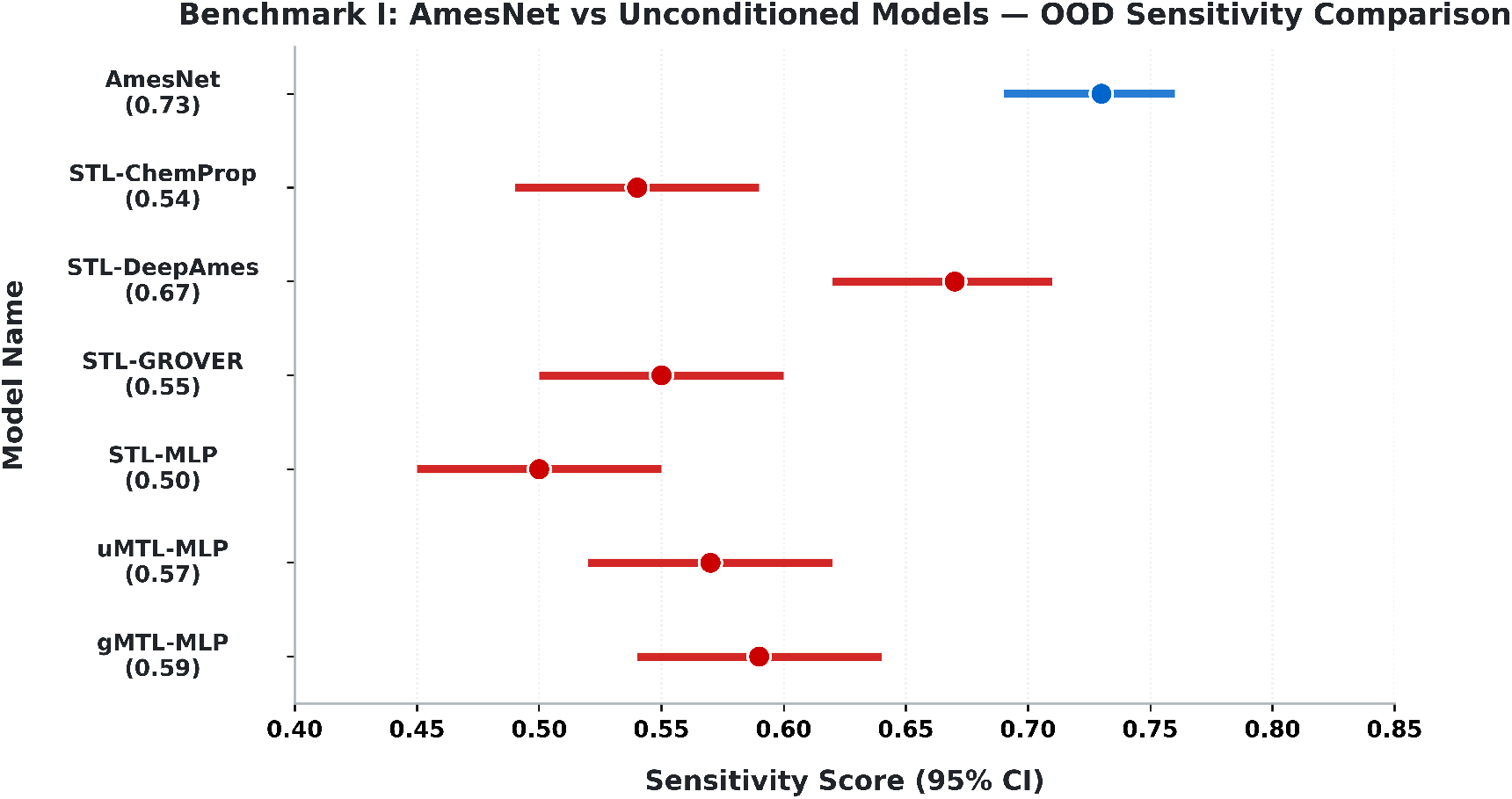
Sample-size-weighted sensitivity on the fixed OOD test set. Error bars indicate 95% confidence intervals from a within-task stratified bootstrap (n = 1,000) across the 16 Ames tasks.

#### Benchmark I: Balanced Accuracy on OOD Test Set

OOD balanced accuracy was calculated with 95% confidence intervals (Figure 4). AmesNet achieved a balanced accuracy of 0.81 (95% CI: 0.79–0.83). AmesNet demonstrates a substantial improvement in balanced accuracy over all models, as evidenced by non-overlapping 95% bootstrap confidence intervals. Within the STL framework, the STL-MLP achieved 0.72 (95% CI: 0.70–0.75), STL-ChemProp achieved (95% CI: 0.72–0.77), STL-GROVER achieved 0.75 (95% CI: 0.73–0.78), and STL-DeepAmes achieved 0.75 (95% CI: 0.72–0.77). Within the uMTL framework, the uMTL-MLP achieved 0.73 (95% CI: 0.70– 0.75). Within the gMTL framework, the gMTL-MLP achieved 0.75 (95% CI: 0.73–0.77). AmesNet achieved balanced accuracy improvements of +0.09 (13%) over STL-MLP, +0.07 (9%) over STL-ChemProp, +0.06 (8%) over STL-GROVER, +0.06 (8%) over STL-DeepAmes, +0.08 (11%) over uMTL-MLP, and +0.06 (8%) over gMTL-MLP.

**Figure 4.**
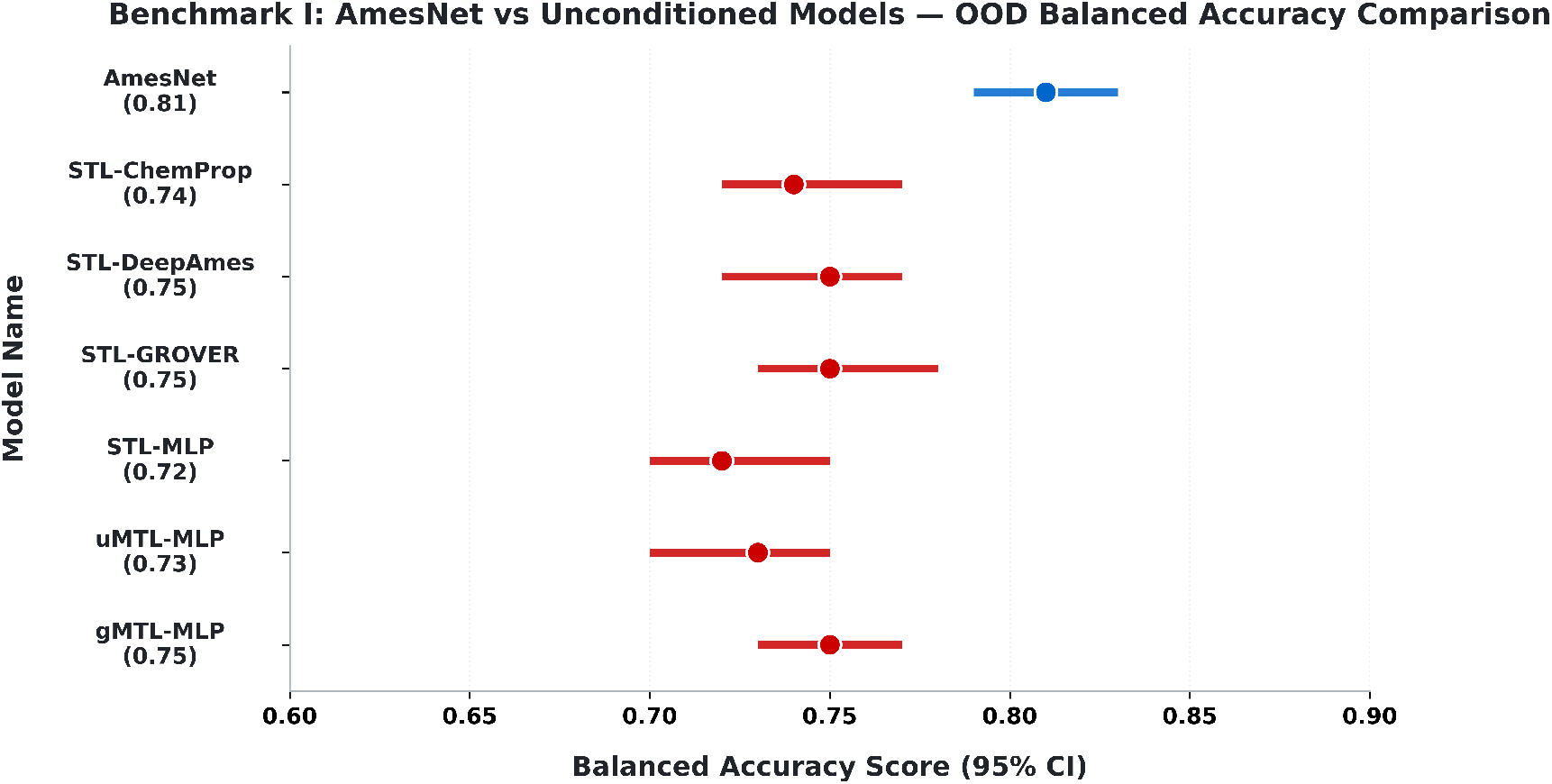
Sample-size-weighted balanced accuracy on the fixed OOD test set. Error bars indicate 95% confidence intervals from a within-task stratified bootstrap (n = 1,000) across the 16 Ames tasks.

#### Benchmark I: Confusion Matrix on OOD Test Set

Figure 5 reports OOD test set confusion matrices. Compared with each baseline, AmesNet increased true positives and decreased false negatives by 123 (STL-MLP), 96 (STL-ChemProp), 91 (STL-GROVER), 27 (STL-DeepAmes), 85 (uMTL-MLP), and 76 (gMTL-MLP).

**Figure 5.**
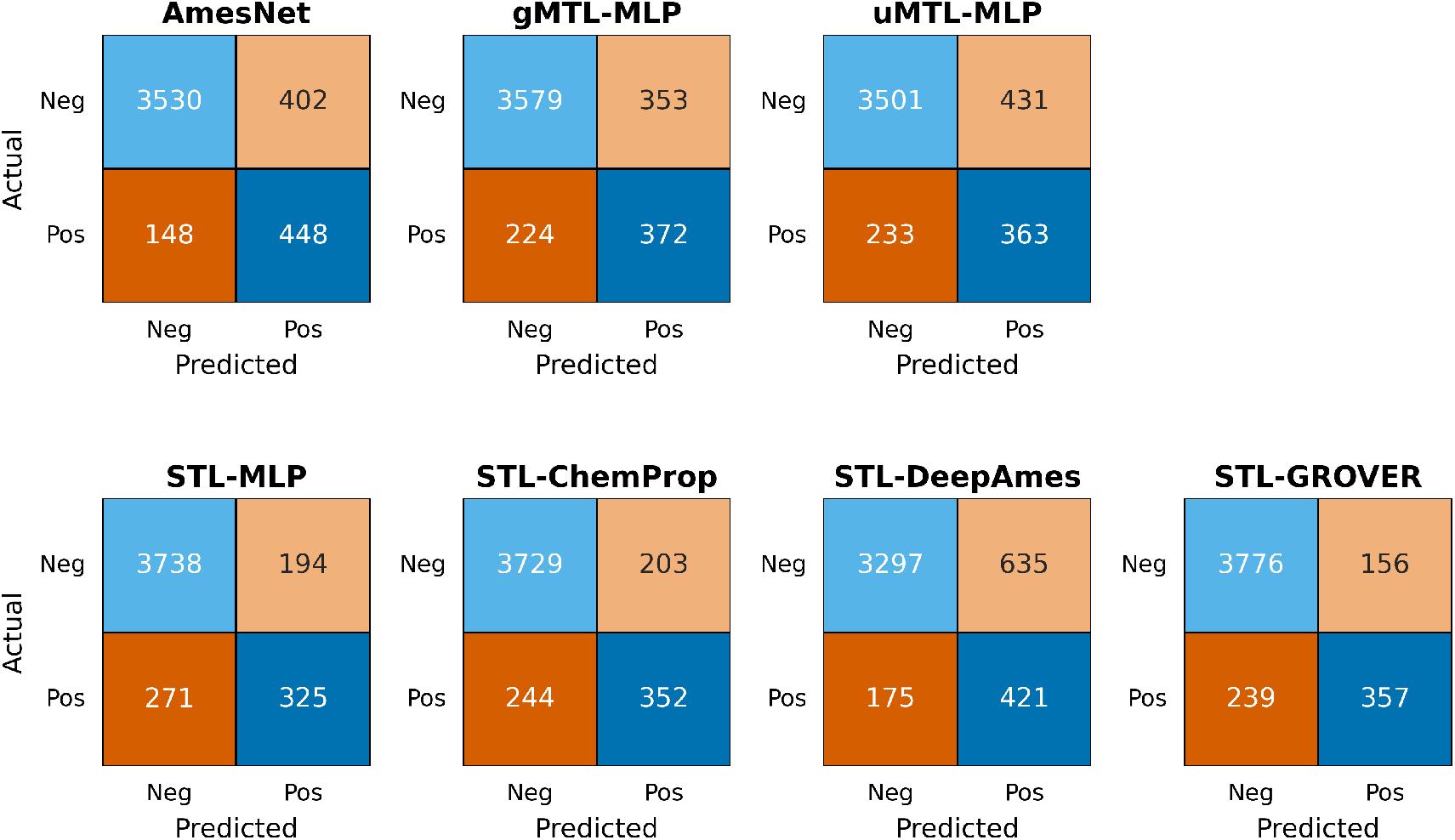
Confusion matrices for AmesNet, gMTL-MLP, uMTL-MLP, STL-MLP, STL-ChemProp, STL-DeepAmes, and STL-GROVER on OOD mutagenicity predictions (n=4,528), showing true negatives, false positives, true positives, and false negatives (clockwise from top-left).

#### Benchmark I: Structural Enrichment of True-Positive Predictions

Figure 6 summarizes mutagenic substructures present in compounds that AmesNet classified as true positives but that Unconditioned Models misclassified as false negatives. The largest counts were observed for planar aromatic intercalators (n = 27–46), aromatic amines (n = 20–33), and aromatic N-heterocycles (n = 17–25). Smaller but recurrent categories included *α,β*-unsaturated carbonyls (n = 6–15), epoxides (n = 5–14), nitro-aromatics (n = 5–11), aromatic ring N-oxides (n = 5–11), and azo groups (n = 4–6). Several alerts were rare across models such as nitroso groups (n = 1–2). These patterns show that a substantial fraction of OOD false negatives in Unconditioned Models are concentrated in a small set of well-known mutagenic alert classes, and that AmesNet correctly identifies many of these cases.

**Figure 6.**
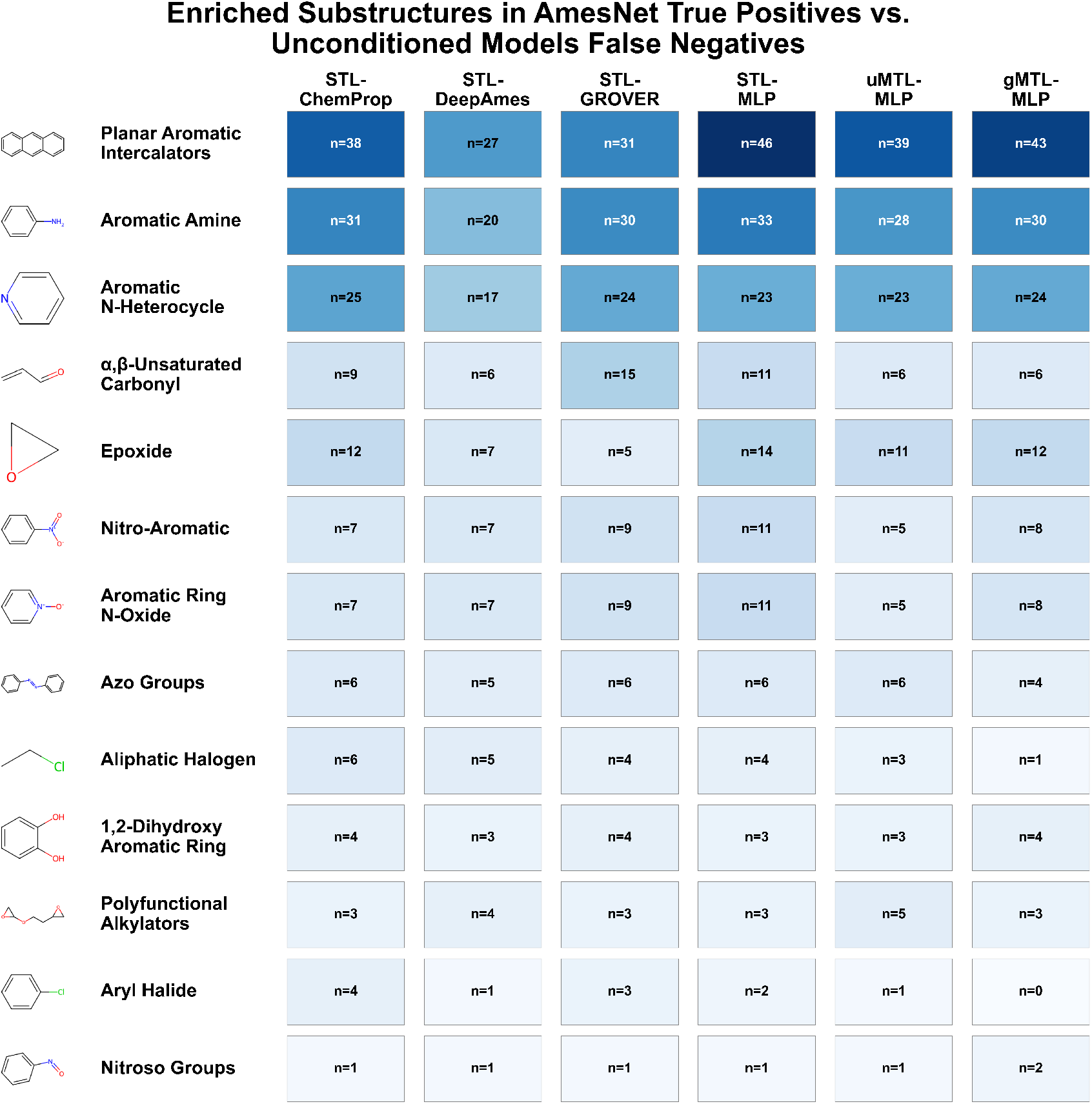
Enriched Substructures in AmesNet True Positives vs. Unconditioned Models False Negatives. Mutagenic substructures observed in compounds correctly predicted by AmesNet but misclassified by unconditioned baseline models (gMTL-MLP, uMTL-MLP, STL-MLP, STL-ChemProp, STL-DeepAmes, STL-GROVER). Values (n) indicate the number of unique substructure-containing compounds where AmesNet achieved true positives while comparison models achieved false negatives.

### Benchmark II: Encoder-Swaps within Task-Conditioning

#### Benchmark II: Sensitivity on OOD Test Set

Benchmark II isolated the contribution of the TCL framework by augmenting two established encoders with the conditioning channel to create TCL-ChemProp and TCL-GROVER. AmesNet achieved the highest sensitivity on the OOD test set (Figure 7), reaching a sensitivity of 0.73 (95% CI: 0.68–0.77). TCL-ChemProp improved sensitivity relative to STL-ChemProp, increasing from 0.54 (95% CI: 0.49–0.59) to 0.65 (95% CI: 0.60–0.70), a +0.11 absolute gain (20% relative). By contrast, TCL-GROVER showed no material sensitivity change relative to STL-GROVER (0.54, 95% CI: 0.50–0.58 vs. 0.55, 95% CI: 0.50–0.60). Together, these results indicate that within the TCL paradigm, AmesNet’s molecular representation provides a sensitivity advantage over leading message-passing and graph transformer alternatives. The TCL framework also confers a material sensitivity lift for TCL-ChemProp relative to STL-ChemProp.

**Figure 7.**
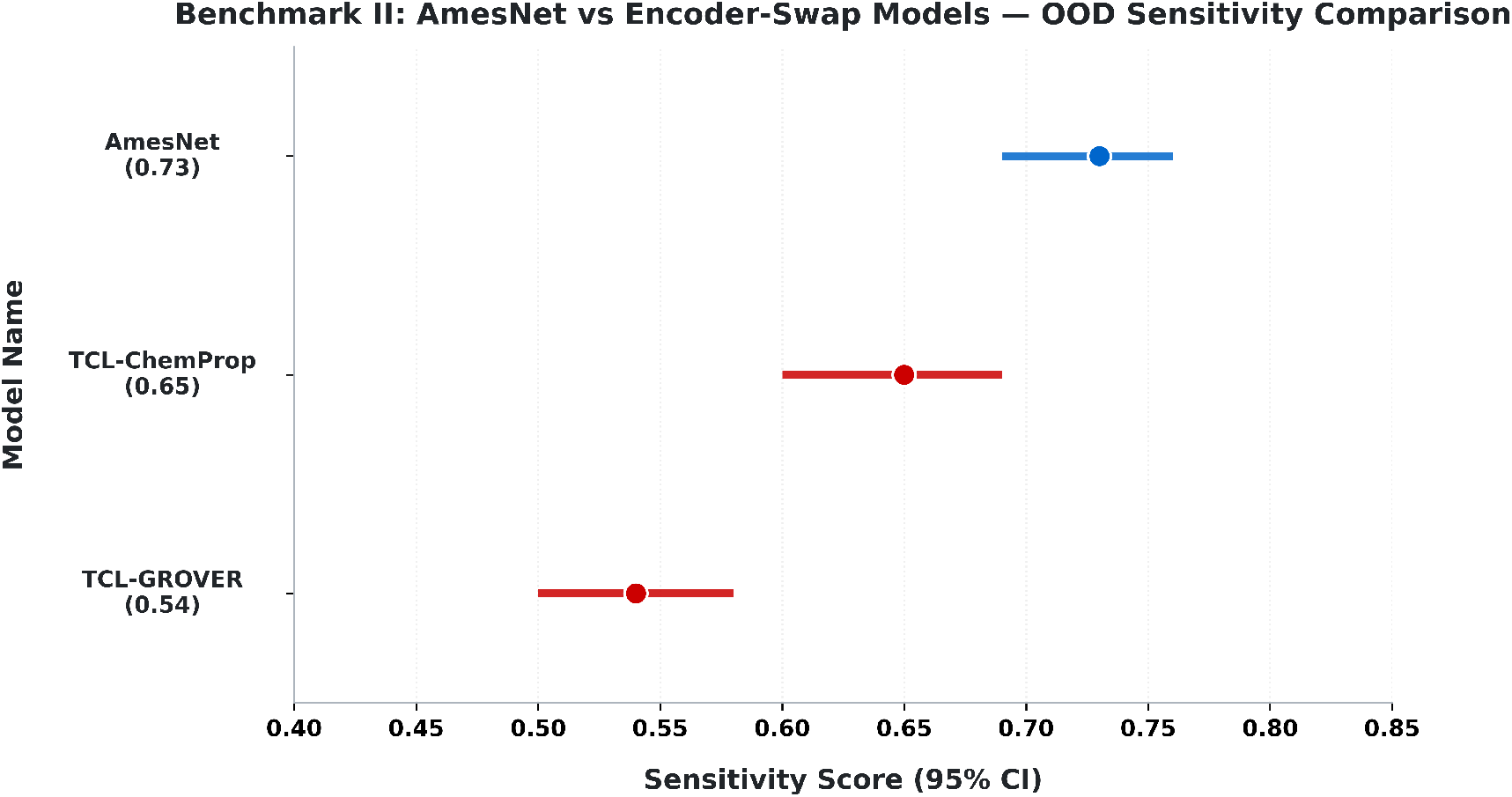
Sample-size-weighted sensitivity on the fixed OOD test set. Error bars indicate 95% confidence intervals from a within-task stratified bootstrap (n = 1,000) across the 16 Ames tasks.

#### Benchmark II: Balanced Accuracy on OOD Test Set

Balanced accuracy on the OOD test set was calculated and 95% confidence intervals were reported (Figure 8). AmesNet achieved a balanced accuracy of 0.81 (95% CI: 0.79–0.83). STL-ChemProp achieved (95% CI: 0.72–0.77) and TCL-ChemProp achieved 0.79 (95% CI: 0.76–0.81). TCL-GROVER showed no material change in balanced accuracy compared to STL-GROVER with 0.75 (95% CI: 0.73–0.77) and (95% CI: 0.73–0.78), respectively. Together, these results show that AmesNet achieves the strongest balanced accuracy and that the TCL framework confers a material balanced accuracy lift for TCL-ChemProp relative to STL-ChemProp.

**Figure 8.**
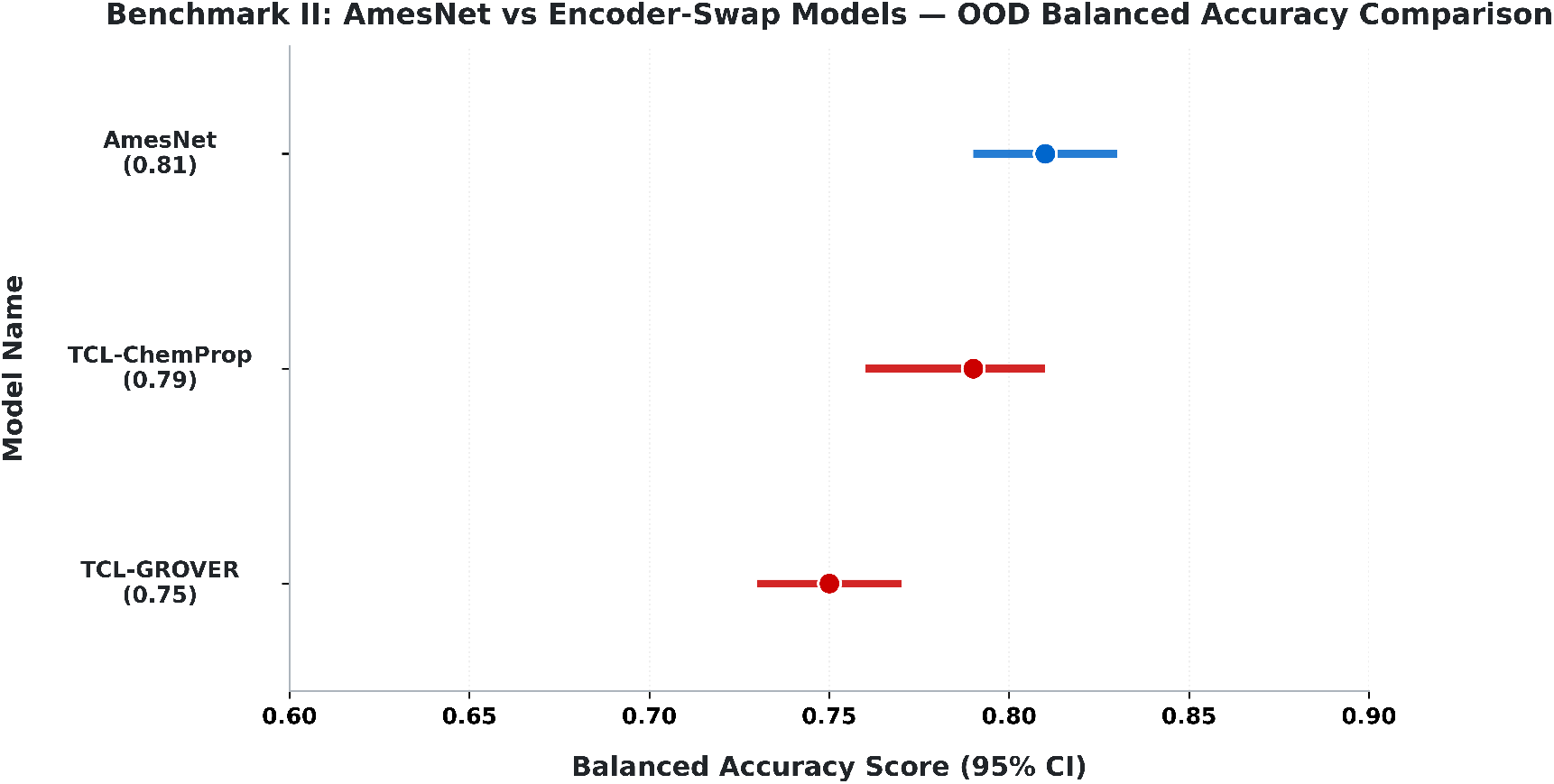
Sample-size-weighted balanced accuracy on the fixed OOD test set. Error bars indicate 95% confidence intervals from a within-task stratified bootstrap (n = 1,000) across the 16 Ames tasks.

#### Benchmark II: Confusion Matrix on OOD Test Set

TCL model confusion matrices on the OOD test set are reported in Figure 9. AmesNet increased true positives and reduced false negatives relative to TCL-ChemProp and TCL-GROVER, demonstrating improved positive-class identification in an OOD setting. AmesNet increased true positives and decreased false negatives by 44 relative to TCL-ChemProp and by 102 relative to TCL-GROVER. TCL-ChemProp increased true positives and decreased false negatives by 52 relative to STL-ChemProp, demonstrating lift associated with the TCL framework. Together, these results indicate that within the TCL paradigm, Ames Net achieves stronger recovery of mutagenic compounds than state-of-the-art message-passing and graph transformer alternatives, and that moving from STL to TCL provides a measurable lift.

**Figure 9.**
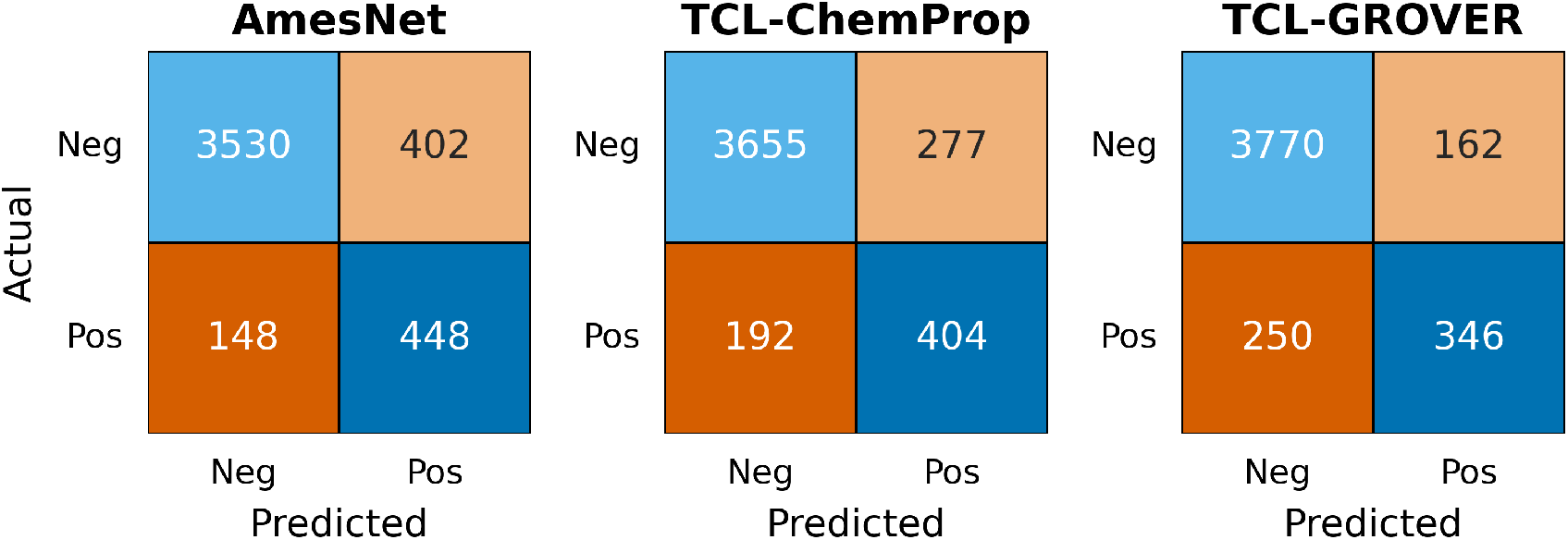
Confusion matrices for AmesNet, TCL-ChemProp, and TCL-GROVER models on OOD mutagenicity predictions (n=4,528), showing true negatives, false positives, true positives, and false negatives (clockwise from top-left).

#### Benchmark II: Structural Enrichment of True-Positive Predictions

Structural enrichment analysis (Figure 10) highlights mutagenic substructures that AmesNet recovers as true positives but that TCL-ChemProp and TCL-GROVER miss as false negatives. The most frequent missed classes were planar aromatic intercalators (n=26 for both models) and aromatic amines (n=24 for TCL-ChemProp; n=26 for TCL-GROVER), followed by aromatic N-heterocycles (n=16–17). Additional recurrent alerts included *α,β*-unsaturated carbonyls (n=6–9), epoxides (n=1–5), nitro-aromatics and aromatic ring N-oxides (n=3–4 each), azo groups (n=5–8), and 1,2-dihydroxy aromatic rings (n=4–7). Relative to STL-ChemProp, TCL-ChemProp shows broad recovery for planar aromatic intercalators (38 to 26), aromatic amines (31 to 24), aromatic N-heterocycles (25 to 16), epoxides (12 to 1), *α,β*-unsaturated carbonyls (9 to 6), nitro-aromatics (7 to 3), and aromatic ring N-oxides (7 to 3). Recovery from STL-GROVER to TCL-GROVER is mixed: TCL-GROVER reduces misses for planar aromatic intercalators (31 to 26), aromatic amines (30 to 26), aromatic N-heterocycles (24 to 17), *α,β*-unsaturated carbonyls (15 to 9), nitro-aromatics (9 to 4), and aromatic ring N-oxides (9 to 4). TCL-GROVER shows no change for epoxides (5 to 5) and increases for azo groups (6 to 8), 1,2-dihydroxy aromatic rings (4 to 7), and nitroso groups (1 to 2). Together, these patterns indicate that within the TCL paradigm, AmesNet’s molecular representation more reliably recognizes mutagenic alert classes than TCL-ChemProp and TCL-GROVER. These results also show that the TCL framework can reduce conserved false-negative failures in established encoders.

**Figure 10.**
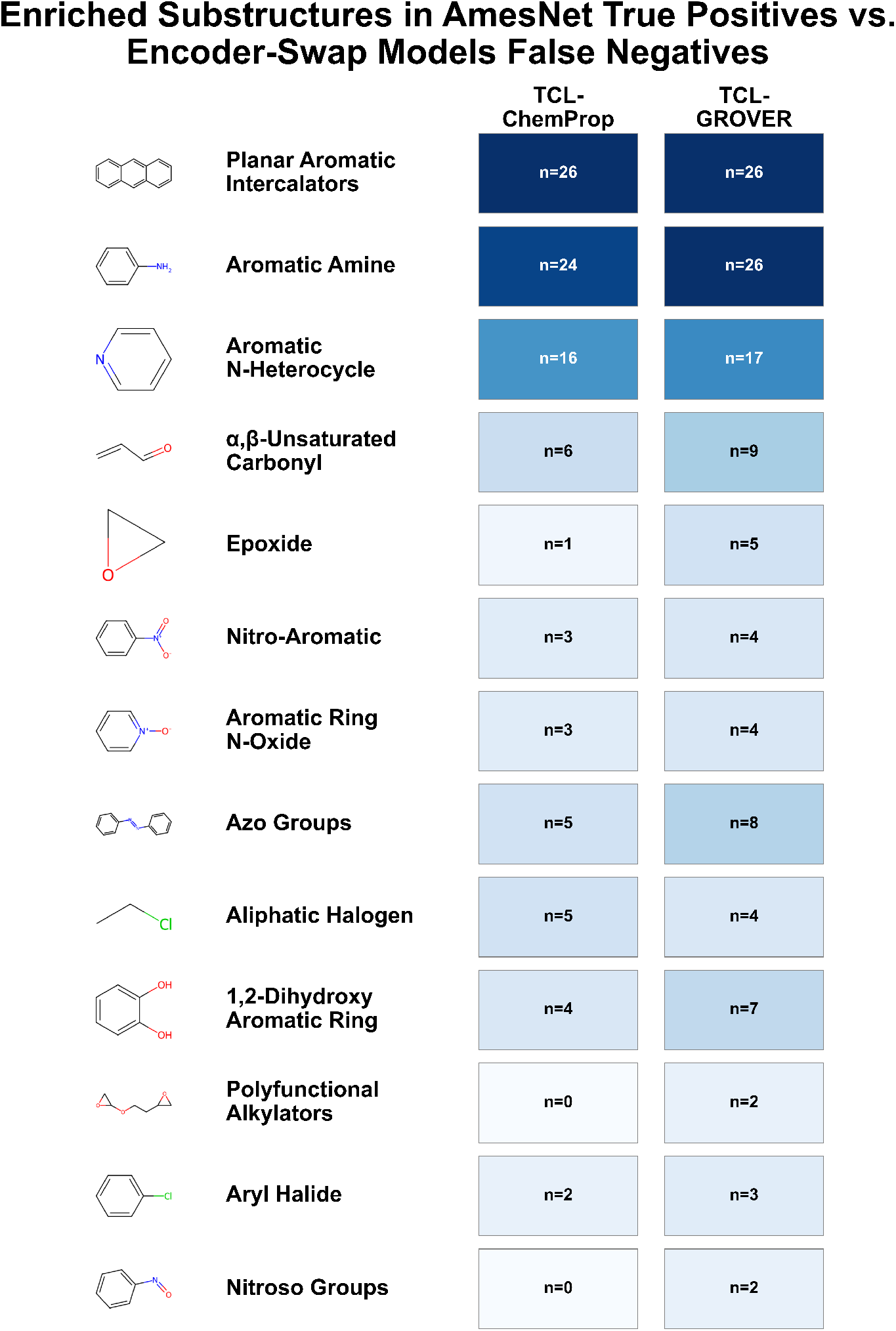
Enriched Substructures in AmesNet True Positives vs. Encoder-Swap TCL Models False Negatives. Mutagenic substructures observed in compounds correctly predicted by AmesNet but misclassified by encoder-swap baseline models (TCL-ChemProp, TCL-GROVER). Values (n) indicate the number of unique substructure-containing compounds where AmesNet achieved true positives while comparison models achieved false negatives.

## Discussion

### Addressing the Performance Gap

#### OOD Sensitivity is a Failure Mode in Ames QSAR

The First and Second Ames/QSAR International Challenge Projects established a consistent failure mode for Ames QSAR models under OOD evaluation. Sensitivity fails to hold under chemical novelty assessment as demonstrated by the participant average sensitivity of 0.46. ChemProp and DeepAmes yielded sensitivities of 0.32 and 0.47 in that competition setting. Those competition results show that existing models do not deliver the sensitivity required for reliable safety screening in chemically novel space.

#### Benchmark I: AmesNet Performance vs. Unconditioned Modeling

We target this failure mode directly using an OOD benchmark. AmesNet shows that sensitivity drop-off is not inevitable under chemical novelty. Instead, this pattern reflects the limitations of the Unconditioned Modeling paradigm that has defined all prior approaches. AmesNet achieves a sensitivity of 0.73 (95% CI: 0.68–0.77), exceeding all Unconditioned Model comparators (Figure 3). AmesNet exhibits non-overlapping 95% confidence intervals against all models except DeepAmes, which still has a lower point-estimate sensitivity. Importantly, the sensitivity improvement of AmesNet is not achieved through a tradeoff in overall discrimination. Balanced accuracy and confusion-matrix analysis were used to assess tradeoffs. AmesNet achieved a balanced accuracy of 0.81 (95% CI: 0.79–0.83) and has non-overlapping 95% confidence intervals against all models (Figure 4). AmesNet’s performance advantage over DeepAmes becomes clear once balanced accuracy is also assessed. AmesNet achieves substantial sensitivity gains without sacrificing overall predictive robustness.

#### Benchmark I: Interpreting the Chemistry of False-Negative Recovery

Structural enrichment analysis demonstrates that AmesNet recovers established mutagenic substructures that escape detection under the traditional Unconditioned Modeling paradigm. AmesNet’s true-positive predictions are enriched for the mutagenic alerts reported in Figure 6. These substructures are well-established sources of false negatives in Ames QSAR modeling[22, 23]. All benchmarked Unconditioned Models exhibit systematic difficulty identifying compounds containing these alerts under OOD evaluation. However, the relative burden of false negatives differs across modeling paradigms. Planar aromatic intercalators and many aromatic N-heterocycles often drive mutagenicity through DNA intercalation and are associated with a frameshift mutation signature[24]. These chemotypes are detected in the Ames frameshift-sensitive bacterial strains (e.g TA1537, TA1538, TA97, and TA98) rather than in base-substitution strains. AmesNet’s recovery of these motifs indicates that it captures strain-dependent positive signals that Unconditioned models miss. Similar context dependence arises for metabolic activation. Aromatic amines require metabolic activation (+S9) to express mutagenicity[25]. AmesNet’s recovery of aromatic amines demonstrates that it is capturing Ames positive risk that is dependent on the S9 activation assay context. AmesNet learns different decision rules across assay states rather than averaging them into a single unconditional boundary, providing a plausible mechanistic explanation for the OOD sensitivity gains.

#### Benchmark II: Encoder-Swaps within Task-Conditioning

The contribution of the TCL framework to the overall success of AmesNet was isolated by injecting the same assay-conditioning channel (strain and *±*S9) into two well-established molecular encoders to form TCL-ChemProp and TCL-GROVER. Similarly to AmesNet’s molecular encoder, ChemProp can be classified as a message-passing network encoder, whereas GROVER is classified as a graph transformer architecture. Performance of TCL-ChemProp and TCL-GROVER variants was compared to their STL counterparts to help attribute the lift to the TCL framework (Figures 7–9). TCL-ChemProp exhibits a clear sensitivity lift over STL-ChemProp, increasing from 0.54 (95% CI: 0.49–0.59) to 0.65 (95% CI: 0.60–0.70) with non-overlapping 95% confidence intervals. This sensitivity lift is accomplished without an apparent trade-off in overall performance as supported by an increase in balanced accuracy from 0.74 (95% CI: 0.72–0.77) to 0.79 (95% CI: 0.76–0.81) for STL-ChemProp and TCL-ChemProp, respectively. However, TCL-GROVER shows no material change in performance relative to STL-GROVER. This suggests that the concatenated conditioning channel may couple more effectively with message-passing network embeddings than to transformer-derived embeddings. Another plausible explanation is that GROVER’s pretrained features are less responsive to explicit metadata concatenation. Even with the performance lift from the TCL framework, no model was as performant as AmesNet in both sensitivity and balanced accuracy. This indicates that the success of AmesNet comes both from its TCL framework and its molecular encoder. Together, these results support TCL as a practical mechanism for improving OOD mutagen recovery without an apparent trade-off in overall classification robustness.

#### Benchmark II: Structural Enrichment Under Task-Conditioning

Structural enrichment analysis in Figure 10 shows that TCL produces a clear recovery for TCL-ChemProp compared to STL-ChemProp. TCL-GROVER shows a weaker mixed effect compared to its STL equivalent. AmesNet remains the strongest overall. Relative to STL-ChemProp, TCL-ChemProp reduces conserved false negatives across several major alert classes, including planar aromatic intercalators (38 to 26), aromatic N-heterocycles (25 to 16), and aromatic amines (31 to 24). The shift from STL-GROVER to TCL-GROVER is less consistently favorable, although TCL-GROVER recovers the same planar aromatic intercalators (31 to 26), aromatic amines (30 to 26), aromatic N-heterocycles (24 to 17). This shows that the TCL framework is responsible for the recovery of these mutagenic toxicophores that are dependent on the bacterial strain and S9 activation assay context. Even after these TCL gains, AmesNet still recovers additional mutagenic compounds that both encoder-swap TCL models miss. These AmesNet only recovered false negatives are concentrated in high-impact chemotypes such as planar aromatic intercalators (n=26 in both TCL baselines), aromatic amines (n=24–26), and aromatic N-heterocycles (n=16–17). This supports the conclusion that AmesNet’s advantage reflects both the TCL framework and a molecular representation that more reliably encodes mutagenicity-relevant structure compared to state-of-the-art message-passing and graph transformer architectures. This result establishes the chemical basis for the sensitivity gains achieved through AmesNet and the TCL paradigm.

### Operational Utility

AmesNet provides the operational utility required for high-confidence mutagenicity screening in early drug discovery. Sensitivity is the most critical metric in Ames prediction and false negatives pose the greatest risk to drug development. A model with low sensitivity fails to identify hazardous compounds and allows them to advance through the pipeline undetected. This failure could waste millions in resources and years in development time. AmesNet enables developers to identify these genotoxic liabilities before significant investment occurs. The TCL framework transforms a potential late-stage safety liability into a primary strategic advantage. It provides a high-confidence pathway for screening compound libraries at high-throughput.

Reported sensitivity values can be difficult to interpret without corresponding trade-off metrics. For example, DeepAmes reports a headline sensitivity of 0.87, but the corresponding trade-off performance metrics in DeepAmes’ additional materials show markedly reduced discrimination (specificity = 0.18, MCC = 0.04, balanced accuracy = 0.52). A balanced accuracy of 0.52 means the model is effectively guessing and provides no reliable predictive signal. In practice, DeepAmes would flag the majority of compounds as mutagenic, limiting utility for screening triage. Additionally, DeepAmes’ sensitivity of 0.87 was achieved by sweeping class-weights that maximizes sensitivity on the test set (weight = 16). This reflects post-hoc operating-point selection and is not directly comparable to a standard held-out test evaluation.

### Beyond Ames

The success of AmesNet establishes Task-Conditioned Learning as a superior design paradigm for Ames modeling. AmesNet leverages the OOD generalization capabilities of ChemPrint. This encoder has achieved on-target hit rates up to 100% and selectivity improvements exceeding 15,000-fold in oncology and antiviral programs. These results demonstrate that ChemPrint performs diverse functions within a single pipeline. Ongoing work will extend this TCL framework into a suite of models for additional ADMET endpoints used in preclinical development.

## Conclusions

Toxicity assessment remains a prerequisite for advancing novel small-molecule therapeutics to human trials. The Ames test is a core genotoxicity assay for identifying mutagenic risk. Developers often wait to complete Ames studies until a candidate approaches regulatory submission because these experiments are costly and frequently deprioritized relative to bioactivity optimization. This timing concentrates risk late in development, where an Ames failure can jeopardize >$10 million in capital and multiple years of progress per candidate.

In response, the FDA and international regulatory agencies have issued a clear call to action to operationalize credible *in silico* toxicology approaches that can reduce these late-stage bottlenecks. *In silico* Ames prediction has not reliably mitigated this bottleneck because existing Ames QSAR models do not sustain high sensitivity under chemical novelty. Optimizing for higher sensitivity has historically forced a trade-off in overall model performance, which can be measured by balanced accuracy. Low sensitivity is a critical failure because false negatives allow mutagenic compounds to advance undetected and recreate the same late-stage bottleneck these *in silico* approaches are intended to prevent.

AmesNet addresses this sensitivity failure by introducing a TCL modeling paradigm that conditions predictions on assay state variables. These states include strain identity and metabolic activation (*±*S9). All prior Ames AI models follow the Unconditioned Modeling paradigm and generate predictions without explicit conditioning on assay state. AmesNet has an assay-state conditioning channel and a ChemPrint-derived molecular encoder. ChemPrint has shown experimental generalization to chemically novel space in Model Medicines’ antiviral and oncology programs. In this work, sensitivity and balanced accuracy are evaluated on a held-out OOD benchmark designed to measure model performance under chemical novelty.

AmesNet achieves a sensitivity of 0.73 (95% CI: 0.68–0.77) with a balanced accuracy of 0.81 (95% CI: 0.79– 0.83), improving sensitivity by up to 46% relative to evaluated Unconditioned Models. AmesNet outperforms the FDA’s DeepAmes in both categories, as the latter achieves a sensitivity of 0.67 (95% CI: 0.62–0.71) and a balanced accuracy of 0.75 (95% CI: 0.72–0.77). By simultaneously improving sensitivity and balanced accuracy, AmesNet successfully overcomes the historical trade-off between sensitivity and overall model performance. Structural enrichment analysis further links these improvements to enhanced recognition of historically challenging chemotypes whose mutagenicity is dependent on specific assay conditions like strain and metabolic activation.

The performance lift provided by the TCL framework was evaluated by injecting the assay-conditioning channel into state-of-the-art message-passing (ChemProp) and graph-transformer (GROVER) architectures. TCL simultaneously improves sensitivity and balanced accuracy when injected into message-passing architectures. Results show that AmesNet still outperforms these TCL adjusted state-of-the-art models, indicating that AmesNet’s encoder provides an additional contribution. These results support TCL modeling as a practical advance for reliable *in silico* mutagenicity screening and position AmesNet as a high-confidence early screening tool to reduce late-stage safety attrition risk in modern drug discovery pipelines.

## Limitations

AmesNet was trained and tested on one public dataset that may not capture the full diversity of Ames positive and negative chemistry. While it outperforms existing models, further validation on additional benchmark data is needed. AmesNet focuses solely on mutagenicity and does not yet address other safety endpoints in a full profile preclinical assessment.

## Associated Content

## Data Availability Statement

All data and code (STL, uMTL, gMTL, TCL encoder swaps, and bootstrapping) are available in the following GitHub repository: https://github.com/Model-Medicines/TCL-Ames

## Supporting Information

The Supporting Information is available free of charge.

- CSV files containing the datasets used to develop and evaluate the models (ZIP).
- CSV files of test-set prediction scores for each evaluated model (ZIP).
- Bootstrap performance metric report (sensitivity, specificity, balanced accuracy, MCC) with 95% confidence intervals for all evaluated models (XLSX).

## Funding

This work was not supported by any external funding.

## Notes

### Competing Interest Statement

The authors have declared no competing interest.

### Summary of Updates

This version sharpens the core contribution (Task-Conditioned Ames prediction), restructures the experiments to cleanly attribute gains via a two-tier benchmark, increases methodological transparency (published code), strengthens evidence (bootstrap CIs, confusion-matrix tradeoffs, and structural enrichment analysis), and clarifies operational framing, and limitations.

